# Activation of the adhesion GPCR GPR133 (ADGRD1) by antibodies targeting the N-terminus

**DOI:** 10.1101/2021.09.13.460139

**Authors:** Gabriele Stephan, Joshua D. Frenster, Ines Liebscher, Dimitris G. Placantonakis

**Affiliations:** Department of Neurosurgery, NYU Grossman School of Medicine, New York, NY 10016, USA; Rudolf Schönheimer Institute for Biochemistry, Molecular Biochemistry, University of Leipzig, Leipzig, Germany; Kimmel Center for Stem Cell Biology, NYU Grossman School of Medicine, New York, NY 10016, USA; Laura and Isaac Perlmutter Cancer Center, NYU Grossman School of Medicine, New York, NY 10016, USA; Brain and Spine Tumor Center, NYU Grossman School of Medicine, New York, NY 10016, USA; Neuroscience Institute, NYU Grossman School of Medicine, New York, NY 10016, USA

**Author notes:** **Corresponding authors:** Dimitris G. Placantonakis, MD, PhD, Department of Neurosurgery, 530 First Avenue, Skirball 8R, NYU Grossman School of Medicine, New York, NY 10016, **And** Gabriele Stephan, PhD, Department of Neurosurgery, 550 First Avenue, Smilow 1306, NYU Grossman School of Medicine, New York, NY 10016.

**Keywords:** adhesion G protein-coupled receptor, antibody, signaling, dissociation

## Abstract

We recently demonstrated that GPR133 (ADGRD1), an adhesion G protein-coupled receptor (aGPCR) whose canonical signaling raises cytosolic cAMP, is necessary for growth of glioblastoma (GBM) and is *de novo* expressed in GBM relative to normal brain tissue. We showed that dissociation of autoproteolytically generated N-terminal and C-terminal fragments (NTF and CTF) of GPR133 at the plasma membrane promotes receptor activation and increases signaling. Toward developing biologics modulating GPR133 function, we tested antibodies against the N-terminus of GPR133 for effects on receptor signaling. Treatment of HEK293T cells overexpressing GPR133 with such antibodies increased cAMP levels in a concentration-dependent manner. Analysis of supernatants following antibody treatment revealed complexes of the antibodies with the autoproteolytically cleaved NTF of GPR133. Cells expressing a cleavage-deficient mutant GPR133 (H543R) did not respond to antibody stimulation, suggesting that the effect is cleavage-dependent. The antibody-mediated stimulation of wild-type GPR133, but not the cleavage-deficient H543R mutant, was reproducible in patient-derived GBM cells. These findings provide a paradigm for modulation of GPR133 function with biologics and support the hypothesis that NTF-CTF dissociation promotes receptor activation and signaling.

## Introduction

Adhesion G protein-coupled receptors (aGPCRs) represent the second largest subfamily within the GPCR superfamily (1,2) and have been implicated in numerous physiological processes and disease mechanisms (3–5). Adhesion GPCRs are structurally characterized by an intracellular C-terminus, a seven transmembrane segment domain and a large extracellular N-terminus (2,6,7). While distinct functional domains within the N-terminus are thought to mediate receptor-specific interactions with adjacent cells or the extracellular matrix (ECM) (2), almost all aGPCRs share a conserved GPCR autoproteolysis-inducing (GAIN) domain within the N-terminus. This domain catalyzes intramolecular autoproteolytic cleavage at the GPCR proteolysis site (GPS) within the N-terminus, resulting in an N-terminal (NTF) and a C-terminal fragment (CTF) (8). A prevalent hypothesis in the field is that binding of ligands from adjacent cells or the ECM to the N-terminus, as well as mechanical stimuli, induce conformational changes or NTF-CTF dissociation (3,9–12). These events, in turn, enable the *Stachel* sequence (9,13–19), a tethered internal agonist peptide sequence immediately distal to the GPS, to activate canonical signaling (3,20). However, the exact activation mechanisms likely differ among members of the aGPCR family and are not well characterized.

Our group recently demonstrated part of the mechanism that mediates activation of GPR133 (ADGRD1), a member of group V of aGPCRs (2) implicated in the pathogenesis of glioblastoma (GBM) (21,22), an aggressive brain malignancy (23). The N-terminus of GPR133, which contains a pentraxin (PTX) domain, undergoes autoproteolytic cleavage almost immediately after protein synthesis (24). However, NTF and CTF stay non-covalently bound to each other until they are trafficked to the plasma membrane, where their dissociation boosts canonical signaling mediated by G_αs_, resulting in activation of adenylate cyclase and elevation in cAMP levels (21,24–27). Our finding that dissociation of NTF and CTF increases canonical signaling is in accordance with the previous observation that the CTF of GPR133, when expressed without the NTF, demonstrates hyperactive signaling relative to its full-length counterpart (9). Collectively, our data suggest that the cleaved but non-covalently associated NTF-CTF dimer is signaling-competent, but its dissociation at the plasma membrane enables full activation of receptor signaling.

Here, we demonstrate that antibodies targeting epitopes outside of the GAIN domain of the N-terminus of GPR133 increase receptor-mediated G_αs_ signaling and cAMP levels. Preventing specific antibody binding by deleting the targeted epitope abolishes the effect. Using biochemical approaches, we provide evidence that antibody binding may enhance dissociation of the NTF from the CTF, thereby leading to increased activation. The antibody-mediated activation is dependent on receptor cleavage, because antibodies fail to modulate signaling of a cleavage-deficient GPR133 mutant (H543R). These findings suggest that GPR133 function can be modulated by antibodies, and likely other biologics as well, which can be used as molecular tools in the study of receptor activation, but also as therapeutic platforms in the context of GBM and possibly other malignancies, where GPR133 plays important roles.

## Results

### Activation of GPR133 signaling with antibodies against its N-terminus

To test whether GPR133 signaling is modulated by antibodies binding to the extracellular N-terminus, we transfected HEK293T cells with GPR133 tagged with an N-terminal hemagglutinin (HA) and a C-terminal FLAG epitope (**Fig. 1A**). Overexpression of tagged GPR133 was verified by western blot analysis of whole cell lysates 48 hours after transfection (**Fig. 1B**). As expected (9,24,27), staining with an anti-FLAG antibody detected the CTF (blue arrow, ~25 kDa), staining with an anti-HA antibody detected bands representing the maturely and immaturely glycosylated NTF (green arrows, ~95/75 kDa), and both antibodies detected small amounts of the full-length uncleaved receptor (red arrows, ~100 – 120 kDa). We used a homogeneous time resolved fluorescence (HTRF)-based assay to quantify cAMP concentrations after expression of GPR133 (**Fig. 1C**). In agreement with previously published data (24,25,27), intracellular cAMP levels increased significantly in HEK293T cells overexpressing GPR133 relative to cells transfected with the empty vector (***p<0.001, t-test).

**Figure 1:**
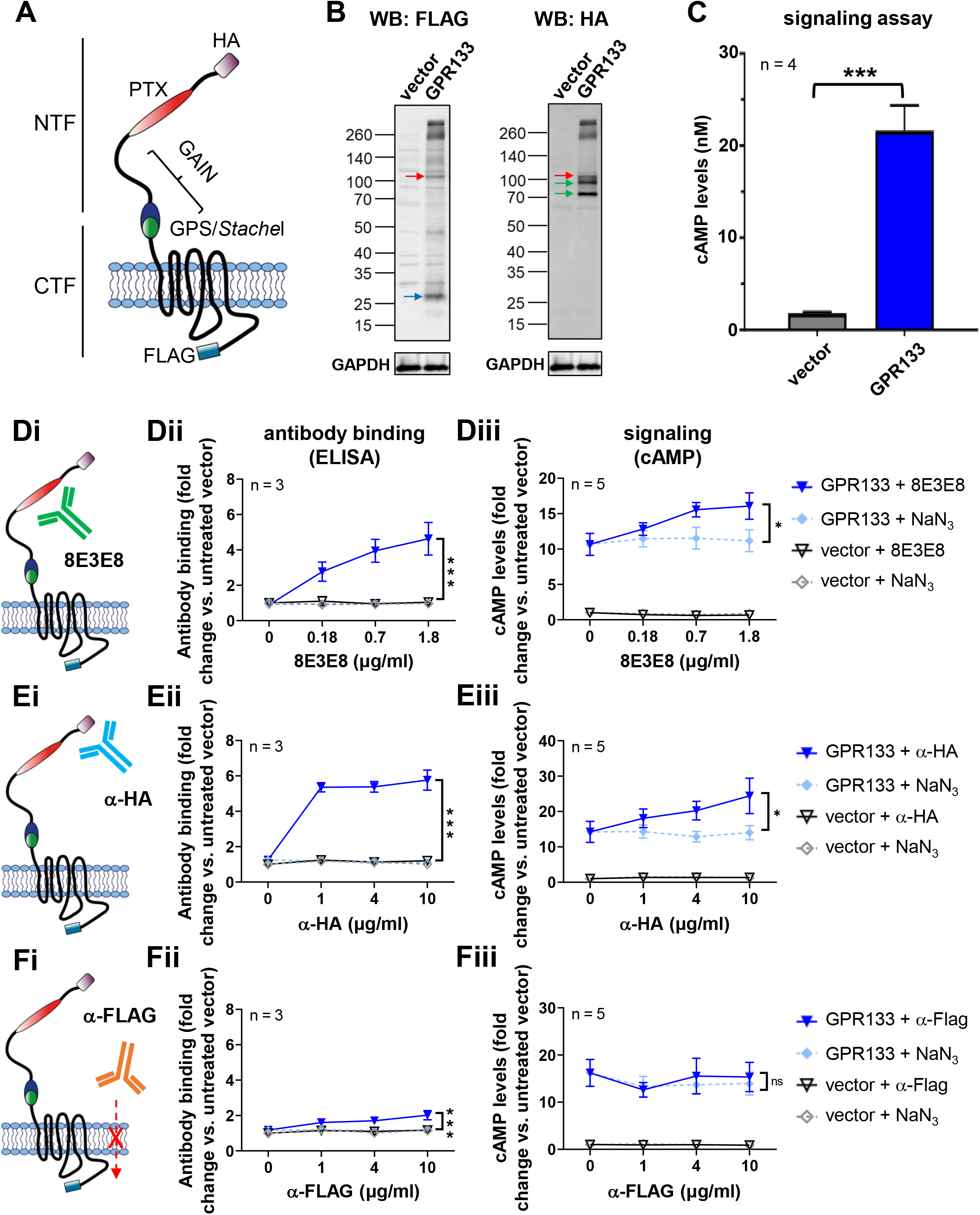
Antibody stimulation increases cAMP levels in HEK293T cells overexpressing GPR133. (**A**) A double-tagged GPR133 construct, including an N-terminal HA-tag and a C-terminal FLAG-tag, was used in these experiments. (**B**) Overexpression of GPR133 is shown by western blots of whole cell lysates. The western blot was simultaneously probed with an anti-HA antibody (targeting the N-terminal HA-tag) and an anti-FLAG antibody (targeting the C-terminal FLAG-tag). The full-length receptor (red arrow), the CTF (blue arrow) and the NTF (green arrow) are detected in cells transfected with GPR133. (**C**) cAMP levels significantly increase after overexpression of GPR133. Bars represent mean ± standard error of the mean (SEM) of 4 individual experiments, ***p<0.001, t-test. (**D-F**) Antibody binding, assessed by ELISA and cAMP levels following treatment of HEK293T cells overexpressing GPR133 with different antibodies. NaN_3_ (0.015 mM, 0.06 mM, 0.15 mM) served as solvent control. Data points represent mean ± SEM of 3-5 individual experiments. The GPR133 + NaN_3_ and GPR133 + 8E3E8 (or α-HA, or α-FLAG) groups were compared by two-way ANOVA (*p<0.05; **p<0.01; ***p<0.001). (**Di**) 8E3E8 targets the N-terminal PTX domain, and (**Dii**) binds to GPR133 in a dose-dependent manner in ELISA assays (F_(1,16)_=41.31,***p<0.0001). (**Diii**) Treatment with 8E3E8 leads to a concentration-dependent increase of cAMP levels compared to the NaN_3_ treatment when overexpressing GPR133 (F_(1,32)_ = 6.509, *p=0.0157). (**Ei**) The anti-HA (α-HA) antibody targets an N-terminal HA tag and (**Eii**) shows binding to GPR133-expressing cells on ELISA assays (F_(1,16)_=284.6, ***p<0.0001). (**Eiii**) Treatment with anti-HA antibody significantly increases cAMP levels compared to the NaN_3_ control (F_(1,32)_= 6.997; *p=0.0125). (**Fi**) HEK293T cells overexpressing GPR133 were treated with an anti-FLAG (α-FLAG) antibody targeting the intracellular C-terminus. (**Fii**) Binding of α-FLAG to GPR133 was statistically significant (F_(1,16)_=17.09, ***p=0.0008) but much less prominent than binding of 8E3E8 or □-HA, and (**Fiii**) there were no significant changes in cAMP concentrations.

We then treated the HEK293T cells with either a mouse monoclonal antibody (8E3E8) that we raised against the PTX domain of GPR133 (**Fig. 1Di**) (21,28), a commercial anti-HA antibody (**Fig. 1Ei**), or a commercial anti-FLAG antibody (**Fig. 1Fi**). A range of antibody concentrations was tested. Since these antibodies were stored in solution containing NaN_3_, increasing concentrations of NaN_3_ (0.015 mM, 0.06 mM, 0.15 mM) served as control. To verify binding of the antibodies to GPR133, we performed an ELISA under non-permeabilizing conditions (**Fig. 1Dii, Eii, Fii**). Optical density increased proportionally with increasing concentrations of the extracellular antibodies (8E3E8: F_(1,16)_=41.31, ***p<0.0001; anti-HA: F_(1,16)_=284.6, ***p<0.0001; two-way ANOVA), compared to the NaN_3_ control. The anti-FLAG antibody, which recognizes the intracellular FLAG epitope, also showed a slight concentration-dependent increase (F_(1,16)_=17.09, ***p=0.0008; two-way ANOVA); however, the ELISA signal was dramatically smaller than with the 8E3E8 and anti-HA antibodies. The increase in optical density after applying the FLAG antibody might be due to partial permeabilization of the plasma membrane during fixing of the cells.

To test the effect of antibodies on GPR133 signaling, we quantified intracellular cAMP levels following stimulation of HEK293T cells transfected with the empty vector or GPR133. We found significant concentration-dependent increases in cAMP following the treatment with 8E3E8 (F_(1,32)_=6.509, *p=0.0157; two-way ANOVA) and anti-HA antibodies (F_(1,32)_=6.997, *p=0.0125; two-way ANOVA), but not the anti-Flag antibody (F_(1,32)_=0.1115, p=0.7407; two-way ANOVA) (**Fig. 1Diii, Eiii, Fiii**). These findings suggest that antibodies targeting the NTF of GPR133 outside of the GAIN domain increase receptor signaling in HEK293T cells.

To ascertain the specificity of the activating effect of 8E3E8, we deleted the PTX domain (amino acids 79 – 276) of GPR133, which contains the epitope that 8E3E8 recognizes. The deletion is predicted to cause a 22 kDa decrease in molecular weight. Overexpression of HA-tagged GPR133 with the PTX deletion (HA-GPR133 ΔPTX) was confirmed by western blot analysis of whole cell lysates (**Fig. 2A**). Staining with the PTX-recognizing 8E3E8 antibody detected HA-GPR133 but not HA-GPR133 ΔPTX, confirming the PTX deletion (**Fig. 2A**). Staining with the anti-HA antibody and a commercial antibody against the cytosolic C-terminus of GPR133 (anti-CTF) demonstrated the expected size shifts of the full-length receptor and the NTF after deletion of the PTX domain (**Fig. 2A**). Importantly, deletion of the PTX domain did not impair receptor cleavage. Immunofluorescent staining of HEK293T cells overexpressing either full length HA-GPR133 or HA-GPR133 ΔPTX with an anti-HA antibody showed similar staining patterns, suggesting the subcellular localization and membrane trafficking of the mutant receptor is not altered (**Fig. 2B**). The baseline levels of cAMP were significantly reduced in HEK293T cells overexpressing HA-GPR133 ΔPTX compared to cells overexpressing HA-GPR133 (F_(2,9)_=4.362; p=0.0474; one-way ANOVA, Tukey’s *post hoc* test for HA-GPR133 compared to HA-GPR133 ΔPTX *p=0.0389) (**Fig. 2Ci**). While treatment of cells overexpressing HA-GPR133 with either 10 μg/mL anti-HA or 1.8 μg/mL 8E3E8 activated receptor signaling equivalently, only treatment with anti-HA but not 8E3E8 increased cAMP levels in cells expressing HA-GPR133 ΔPTX (F_(2,18)_=9.490, p=0.0015 two-way ANOVA; Tukey’s *post hoc* test: HA-GPR133 control vs. HA-GPR133 + α-HA **p=0.0029; HA-GPR133 control vs. HA-GPR133 + 8E3E8 *p=0.0126; HA-GPR133 ΔPTX control vs. HA-GPR133 ΔPTX + α-HA **p=0.0047; HA-GPR133 ΔPTX control vs. HA-GPR133 ΔPTX + 8E3E8 p=0.9558). This finding indicated specificity of the activating effect of 8E3E8 on GPR133 signaling.

**Figure 2:**
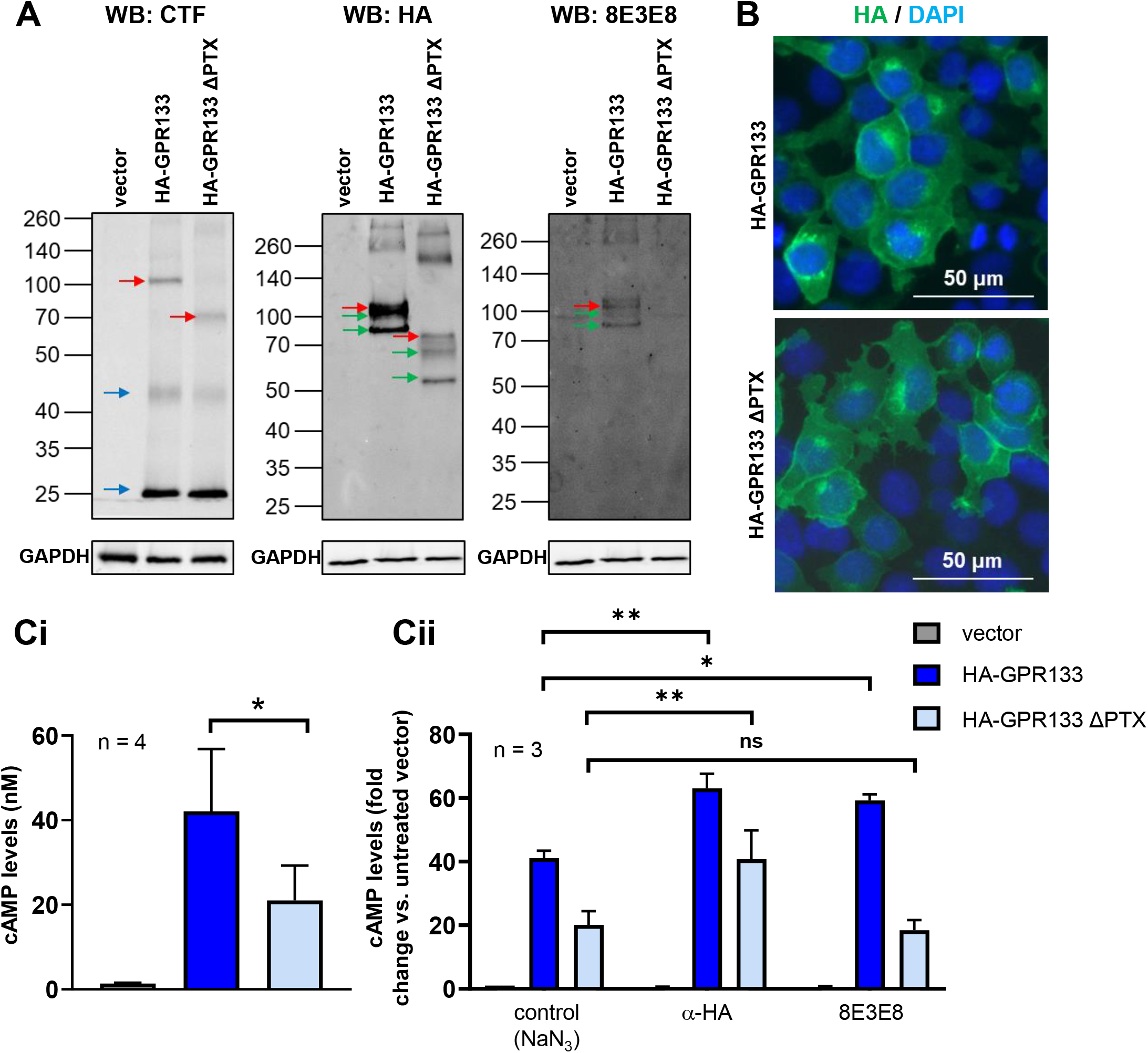
Specific binding of the antibody to the GPR133 PTX domain activates the receptor. (**A**) Western blot analysis of whole cell lysates of HEK293T cells overexpressing HA-GPR133 and the deletion mutant HA-GPR133 ΔPTX, compared to an empty vector. Detection with an HA-antibody and an anti-CTF antibody shows the full-length receptor (red arrow), the NTF (green arrow) and the CTF (blue arrow). Size shifts of the full-length receptor and the NTF are related to the PTX deletion. 8E3E8 detects HA-GPR133, but not the deletion mutant. The same membrane was stained with anti-HA and 8E3E8. (**B**) Immunostaining of HEK293T cells overexpressing HA-GPR133 and HA-GPR133 ΔPTX with an anti-HA antibody (green) and DAPI (blue) shows that both constructs can be detected at the plasma membrane. (**C**) cAMP responses of HEK293T cells overexpressing HA-GPR133 and HA-GPR133 ΔPTX. Bars represent mean ± SEM of 3-4 individual experiments. (**Ci**) Deletion of the PTX domain significantly reduces cAMP levels (F_(2,9)_=4.362; p=0.0474; one-way ANOVA, Tukey’s *post hoc* test for HA-GPR133 compared to HA-GPR133 ΔPTX *p=0.0389). (**Cii**) HA-GPR133 signaling is increased significantly by adding an HA-antibody, binding the N-terminal HA-tag, or 8E3E8, binding the PTX domain. In contrast, HA-GPR133 ΔPTX signaling is increased only by the HA-antibody, but not 8E3E8. (F_(2,18)_=9.490; p=0.0015 two-way ANOVA; Tukey’s *post hoc* test: HA-GPR133 control vs. HA-GPR133 + □-HA **p=0.0029; HA-GPR133 control vs. HA-GPR133 + 8E3E8 *p=0.0126; HA-GPR133 ΔPTX control vs. HA-GPR133 ΔPTX + □-HA **p=0.0047; HA-GPR133 ΔPTX control vs. HA-GPR133 ΔPTX + 8E3E8 p=0.9558).

We then tested the hypothesis that increasing the effective concentration of antibodies at the cell surface would further enhance GPR133 signaling. To accomplish this, we coupled 8E3E8 or the HA-antibody to Dynabeads® (**Fig. 3**) and treated HEK293T cells overexpressing the HA and FLAG-tagged construct of GPR133 (**Fig. 3Ai, Bi**). Unconjugated Dynabeads® served as a control. We visualized the binding interaction between the antibody-coated beads and HEK293T cells overexpressing GPR133 with light microscopy. Dynabeads® localized to the plasma membrane of cells only in the condition where they were coated with anti-HA antibody and cells expressed HA-tagged GPR133 (**Supplementary Fig. 1**). In the absence oft he HA tag in GPR133 or anti-HA antibody coating oft he beads, the Dynabeads appear to cluster in spaces between cells (**Supplementary Fig. 1**). Furthermore, we tracked the beads’ diffusivity over time by microscopic video capture and found that anti-HA antibody-conjugated Dynabeads® had reduced mobility as they bound HA-GPR133 expressing cells, while unconjugated Dynabeads® displayed a higher degree of molecular motion, suggesting they remained unbound to cells (**Supplementary Fig. 2**). Overall, the imaging findings indicated specificity of binding of antibody-coated beads to the cell surface. Accordingly, intracellular cAMP levels increased significantly after treating the cells with increasing concentrations of 8E3E8-conjugated Dynabeads® (**Fig. 3Aii**; F_(1,12)_=82.41, ***p<0.001, two-way ANOVA) or anti-HA-conjugated Dynabeads® (**Fig. 3Bii**; F_(1,12)_=56.46, ***p<0.001, two-way ANOVA) compared to cells treated with unconjugated Dynabeads®. The effect of antibody-coated beads on GPR133 signaling appeared larger than the effect of antibodies alone (compare to **Fig. 1**), suggesting that increased effective concentrations of antibodies at the cell surface promotes receptor activation. Taken together, these findings indicate that antibodies against the N-terminus of GPR133 increase receptor signaling.

**Figure 3:**
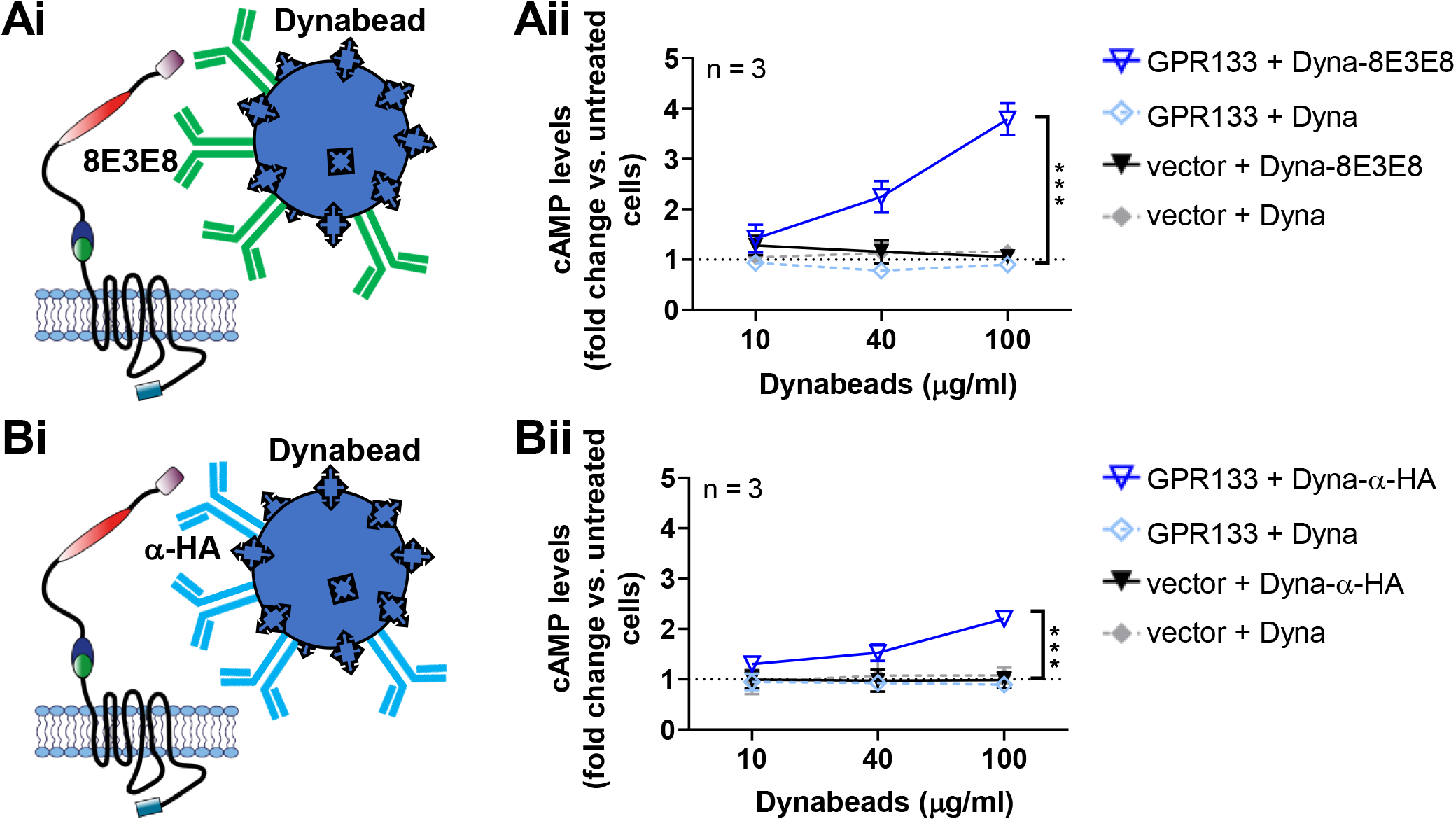
Stimulation with antibody-conjugated Dynabeads® increases cAMP levels in HEK293T cells overexpressing GPR133. HEK293T cells overexpressing a double-tagged GPR133 construct, were treated with Dynabeads® conjugated to (**Ai**) 8E3E8 or (**Bi**) anti-HA antibodies. Data points represent mean ± SEM of 3 individual experiments. (**Aii**) Treatment with 8E3E8-conjugated Dynabeads® leads to a concentration-dependent increase of cAMP levels compared to the treatment with unconjugated beads (F_(1,12)_=82.41, ***p<0.001, two-way ANOVA). (**Bii**). Treatment with anti-HA-conjugated Dynabeads® significantly increases cAMP levels compared to unconjugated beads (F_(1,12)_=56.46, ***p<0.001, two-way ANOVA).

### Antibody binding stimulates dissociation of the GPR133 NTF from the CTF

We recently showed that NTF-CTF dissociation at the plasma membrane boosts GPR133 canonical signaling (24). We therefore hypothesized that the effects of antibodies on GPR133 signaling may be mediated by antibody-induced NTF-CTF dissociation at the plasma membrane (**Fig. 4A**). To test this hypothesis, we transfected HEK293T cells with GPR133 tagged with Twin-Strep-tag at the N-terminus (**Fig. 4A**) and treated them with 1.8 μg/mL 8E3E8, which binds the PTX domain of the N-terminus. Western blot analysis of whole cell lysates using an anti-Strep antibody confirmed that the tagged GPR133 was overexpressed and cleaved as expected (27), with the cleaved NTF (green arrow) being the dominant portion detected (**Fig. 4Bi**). The anti-CTF antibody detected the cleaved CTF (blue arrow), as well as small amounts of the uncleaved full-length receptor (red arrow) (**Fig. 4Bii**).

**Figure 4:**
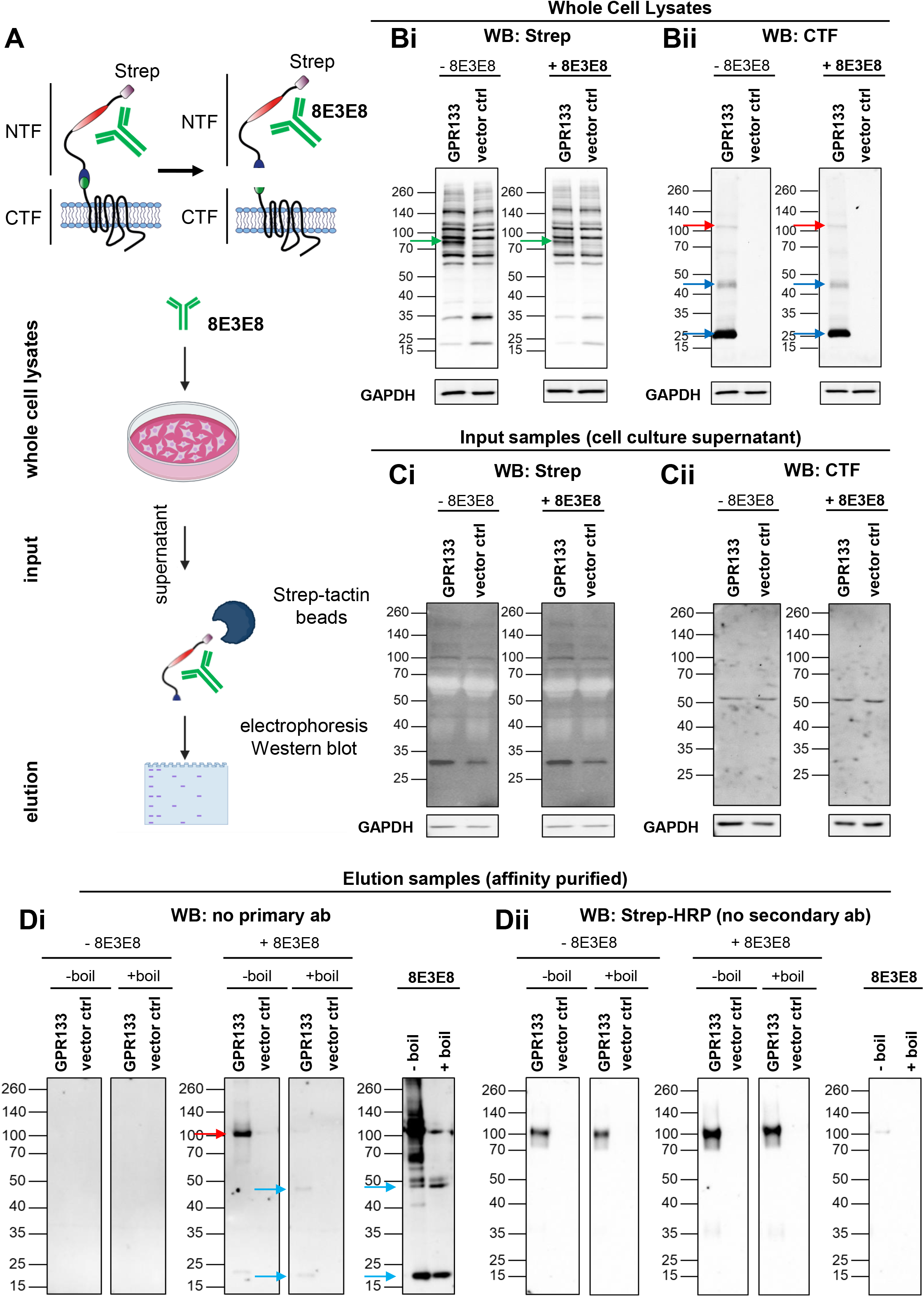
The 8E3E8 antibody is bound to the GPR133 NTF in the supernatant. **(A)** Experimental outline. GPR133 constructs used for the experiments (n=6) carried an N-terminal Strep-tag. (**B**) Representative Western blot of whole cell lysates. (**Bi**) An anti-Strep antibody was used to detect the NTF (green arrow) and (**Bii**) anti-CTF was used to show the CTF (blue arrow) and the uncleaved full-length receptor (red arrow). GAPDH served as loading control. (**C**) Western blot of input samples, representing the supernatant of HEK293T cells overexpressing GPR133 with and without treatment with 8E3E8, probed with an anti-Strep antibody (**Ci**) or anti-CTF (**Cii**). The NTF and CTF are not detectable. (**D**) Western blot analysis of elution samples and the 8E3E8 antibody itself without the use of a primary antibody. Bands were detected using a secondary antibody only, indicating that 8E3E8 is bound to the GPR133 NTF after affinity purification from the supernatant (red arrow). Boiling of elution samples and 8E3E3 itself resolves heavy and light chains of 8E3E8 (blue arrows). (**Dii**) Western blot analysis of boiled and un-boiled elution samples using an anti-Strep antibody conjugated to horseradish peroxidade (HRP), which directly detects the GPR133 NTF and obviates the need for a secondary antibody. The elution samples and 8E3E8 were prepared the same way as in **Figure 4Di**.

We then collected supernatants after the antibody treatment. Western blot analysis of the supernatants with an anti-Strep antibody or an anti-CTF antibody did not show any GPR133-specific bands (**Fig. 4Ci,ii**). To test if the 8E3E8 antibody is still bound to the Twin-Strep-tagged NTF after affinity purification from the cell culture supernatant with Strep-Tactin® XT coated magnetic beads, we developed approaches to identifying each component of the NTF-8E3E8 complex. First, to identify 8E3E8, we probed western blots of supernatant eluates after antibody 8E3E8 treatment with an anti-mouse secondary antibody (anti-IgG1) in the absence of a primary antibody (8E3E8 is a mouse monoclonal IgG1 antibody), and detected a band at ~100 kDa (**Fig. 4Di**, red arrow). To verify that this band represents antibody-NTF complexes that include 8E3E8, we boiled the elution samples and reran the western blot, again probing only with the anti-mouse secondary antibody. Boiling revealed the heavy and light chain of the 8E3E8 antibody (**Fig. 4Di**, blue arrows) and produced a banding pattern similar to that of the boiled 8E3E8 antibody (**Fig. 4Di**).

Second, to directly identify NTF in these presumed complexes, we probed eluates with an anti-Strep antibody (which, like 8E3E8, is a mouse IgG1 antibody) conjugated to HRP, thus forgoing the need for a secondary antibody. Indeed, we were able to detect a GPR133-specific band at ~100 kDa representing the NTF (**Fig. 4Dii**). These findings suggest that 8E3E8 is bound to the NTF in the supernatant, which supports the hypothesis that the activating effect of the antibody on GPR133 signaling is primarily mediated by NTF-CTF dissociation.

Finally, we analyzed the affinity-purified Strep-tagged NTF of GPR133 by western blot using the HRP-conjugated anti-Strep antibody (**Fig. 5A**). The densitometric intensity of this protein band was higher in our 8E3E8-treated samples compared to control untreated samples in 3 of 6 individual experiments (**Fig. 5B**). This finding suggested that binding of 8E3E8 increased the dissociation of the NTF from the CTF. The fact that this phenomenon was observed in only half of our experiments may be attributable to technical reasons, given that the affinity purification is focused on enrichment of the NTF from the supernatant through the Twin-Strep-Tag.

**Figure 5:**
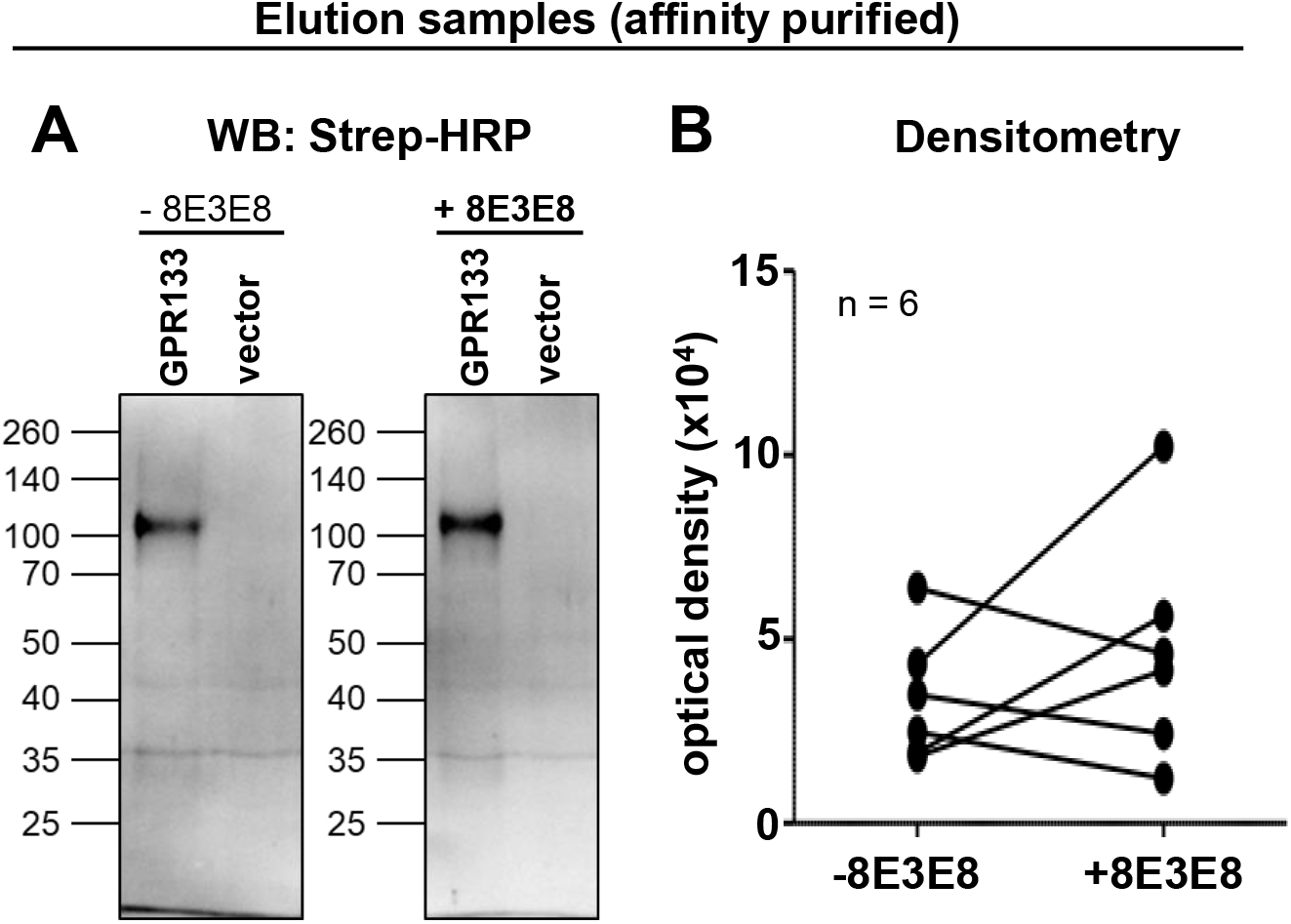
Antibody binding increases dissociation of the cleaved NTF from the CTF. (**A**) Elution of supernatant samples with and without treatment with 8E3E8 after Streptactin® purification of TwinStrep-tagged GPR133. Using an anti-Strep antibody, a protein band representing the NTF of GPR133 at ~100 kDa is detected. (**B**) Densitometry analysis of the elution band shown in (**A**). Data points represent six individual experiments. The intensity of the band, representing the NTF increased in 6 out 3 individual experiments.

To ensure that the protein band detected at ~100 kDa in **Fig. 5A** represents the cleaved NTF of GPR133 and not the uncleaved form of the full-length receptor, we probed eluates with the anti-CTF antibody and did not detect any GPR133-specific bands before or after treatment with 8E3E8 (**Suppl. Fig. 3A**). In an additional quality control experiment, elution samples from the supernatant before and after treatment with 8E3E8 were deglycosylated prior to western blot analysis. Consistent with our prior observations (24), we observed a shift of the apparent molecular weight of the band detected with the anti-Strep antibody from ~100 kDa to ~65 kDa (**Suppl. Fig. 3B**), indicating that the detected band in **Fig. 5A** indeed represents the maturely glycosylated NTF of GPR133.

### Activation of GPR133 by antibodies depends on receptor cleavage

A prediction of the model that antibody-mediated stimulation of GPR133 signaling relies on NTF-CTF dissociation is that the effect should not occur in uncleavable GPR133 mutants. To test this prediction, we overexpressed a cleavage-deficient point mutant (H543R) GPR133 (24,25) with an N-terminal HA tag and a C-terminal FLAG tag (**Fig. 6A**) in HEK293T cells. Overexpression of WT GPR133 and the uncleaved receptor was verified by western blot analysis of whole cell lysates using the anti-HA and anti-FLAG antibodies. Expression of both constructs were similar 48 h after transfection (**Fig. 6B**). Following overexpression of the uncleavable mutant receptor, we detected maturely (black arrow, ~120 kDa) and immaturely glycosylated (red arrow, ~100 kDa), full-length H543R GPR133. In contrast to the WT receptor, bands representing the cleaved NTF or CTF were not detectable, indicating lack of cleavage of the mutant receptor. HTRF analysis showed significantly higher cAMP levels after overexpression of H543R compared to the empty vector control (***p<0.001, t-test) (**Fig. 6C**). We then treated HEK293T cells overexpressing H543R with the same antibodies used to stimulate WT GPR133 (8E3E8, anti-HA and anti-FLAG) (**Fig. 6D-F**), in incrementally increasing concentrations. NaN_3_ served as solvent control. To verify binding of the antibodies to GPR133, we performed an ELISA under non-permeabilizing conditions (**Fig. 6Dii, Eii, Fii**) and found concentration-dependent binding of the extracellular antibodies (8E3E8: F_(1,16)_=54.96, ***p<0.0001; anti-HA: F_(1,16)_=130.1, ***p<0.0001; two-way ANOVA) to H543R GPR133 (**Fig. 6Dii, Eii**). Similar to our experiments with WT GPR133, optical density after treatment with the intracellular FLAG antibody was significantly increased compared to the NaN_3_ control (F_(1,16)_=19.77, ***p=0.0004; two-way ANOVA). However, the magnitude of the ELISA signal was much lower compared to the treatment with 8E3E8 or anti-HA.

**Figure 6:**
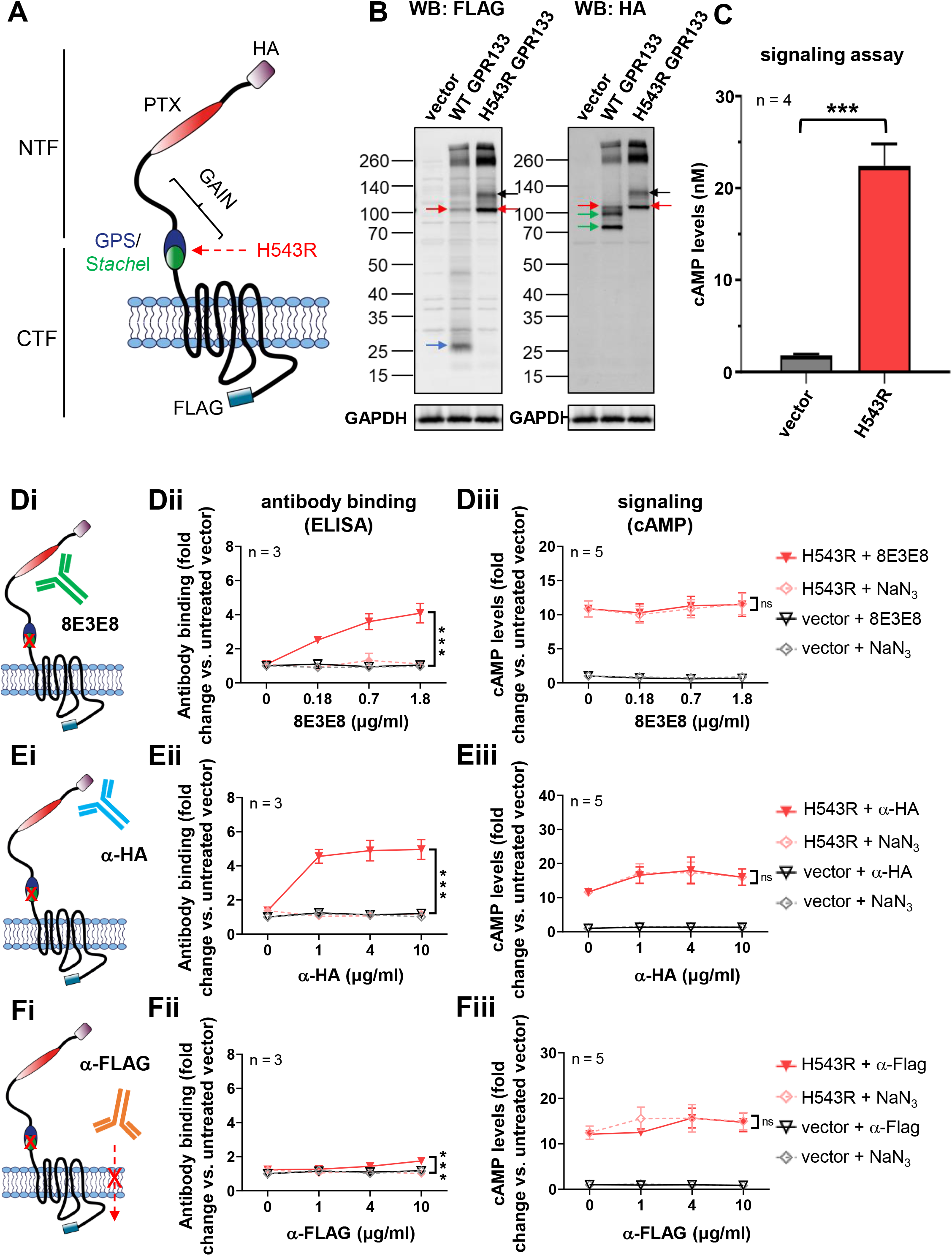
The cleavage-deficient mutant H543R is not responsive to the antibody stimulus. (**A**) The double-tagged GPR133 construct, including an N-terminal HA-tag and a C-terminal FLAG-tag, was mutated at position H543R to prevent receptor cleavage. (**B**) Overexpression of the WT receptor and the cleavage-deficient mutant H453R is shown by western blots of whole cell lysates. The western blot was simultaneously probed with an anti-HA antibody (targeting the N-terminal HA-tag) and an anti-FLAG antibody (targeting the C-terminal FLAG-tag). Overexpression of WT GPR133 shows bands, representing immaturely glycosylated full-length receptor (red arrow), the CTF (blue arrow) as well as the immaturely and maturely glycosylated NTF (green arrow). Overexpression of H543R shows two bands, representing the maturely (black arrow) and immaturely (red arrow) glycosylated full-length receptor. The WT receptor and the cleavage-deficient mutant show similar expression levels. (**C**) cAMP levels increase after overexpression of H543R. Bars represent mean ± SEM of 4 individual experiments. ***p<0.001, t-test. (**D-F**) Antibody binding, assessed by ELISA and cAMP levels following treatment of HEK293T cells overexpressing H543R with different antibodies. NaN_3_ (0.015 mM, 0.06 mM, 0.15 mM) served as solvent control. Data points represent mean ± SEM of 3-5 individual experiments. Statistical significance was assessed by two-way ANOVA (*p<0.05; **p<0.01; ***p<0.001). (**D**) 8E3E8 targeting the PTX domain (**Di**) binds to H543R in a dose-dependent manner on ELISA assays (F_(1,16)_=54.96, ***p<0.0001) (**Dii**). Treatment with 8E3E8 does not lead to a concentration-dependent increase of cAMP levels compared to the NaN_3_ treatment (**Diii**). (**E**) The anti-HA antibody binds the N-terminal HA tag (**Ei**) in dose-dependent fashion by ELISA (F_(1,16)_=130.1, ***p<0.0001) (**Eii**). Treatment with anti-HA did not increase cAMP levels compared to the NaN_3_ control (**Eiii**). (**F**) Treating HEK293T cells overexpressing H543R with a C-terminal anti-FLAG antibody (**Fi**) led to less antibody binding (F_(1,16)_=19.77, ***p=0.0004) (**Fii**) and no significant changes of cAMP levels (**Fiii**).

Next, we quantified intracellular cAMP levels following stimulation with the antibodies (**Fig. 6Diii, Eiii, Fiii**). In contrast to HEK293T cells overexpressing the WT receptor, treatment of cells expressing H543R with 8E3E8 or anti-HA antibodies did not increase cAMP levels compared to the treatment with NaN_3_ (**Fig. 6Diii, Eiii**). As a control, we repeated the assay with the anti-FLAG antibody, recognizing the receptor’s C-terminus, which did not activate the mutant receptor (**Fig. 6Fiii**). These findings suggest that stimulation of receptor signaling by 8E3E8 or anti-HA antibodies is cleavage-dependent.

### Antibody-mediated stimulation of GPR133 is reproducible in GBM cells

Next, we investigated whether our findings with the WT and the cleavage-deficient H543R mutant GPR133 could be reproduced in patient-derived GBM cells. We transfected a patient-derived GBM culture, GBML137, with untagged versions of WT and the cleavage-deficient H543R mutant GPR133, as described in previous studies (24) (**Table 1**, **Fig. 7A**). Western Blot analysis of whole cell lysates, using anti-CTF and anti-NTF (8E3E8) antibodies, confirmed that both constructs were expressed in GBM cells in comparable amounts (**Fig. 7B**). HTRF-based analysis of intracellular cAMP levels showed a significant increase of cAMP after transfection of WT and H543R GPR133, compared to the empty vector (F(_2,12_)=14.01, ***p=0.0007, one way ANOVA, Tukey’s post hoc test: vector vs. GPR133 *p=0.0006; vector vs. H543R *p=0.0113) (**Fig. 7C**). GBM cells overexpressing WT and H543R GPR133 were then treated with the 8E3E8 antibody (1.8 μg/mL) and an antibody targeting the intracellular C-terminus (2 μg/mL anti-CTF). Treatment with 8E3E8 significantly increased cAMP levels relative to the solvent control NaN_3_ in WT GPR133-expressing cells (F_(2,42)_=33.66, p<0.0001, Tukey’s *post hoc* test: GPR133 + NaN_3_ and GPR133 + 8E3E8 **p=0.0091), but not in cells expressing the uncleaved H543R mutant receptor (**Fig. 7D**). Furthermore, the anti-CTF antibody, which is not expected to permeate the plasma membrane, had no effect on signaling (**Fig. 7D**).

**Figure 7:**
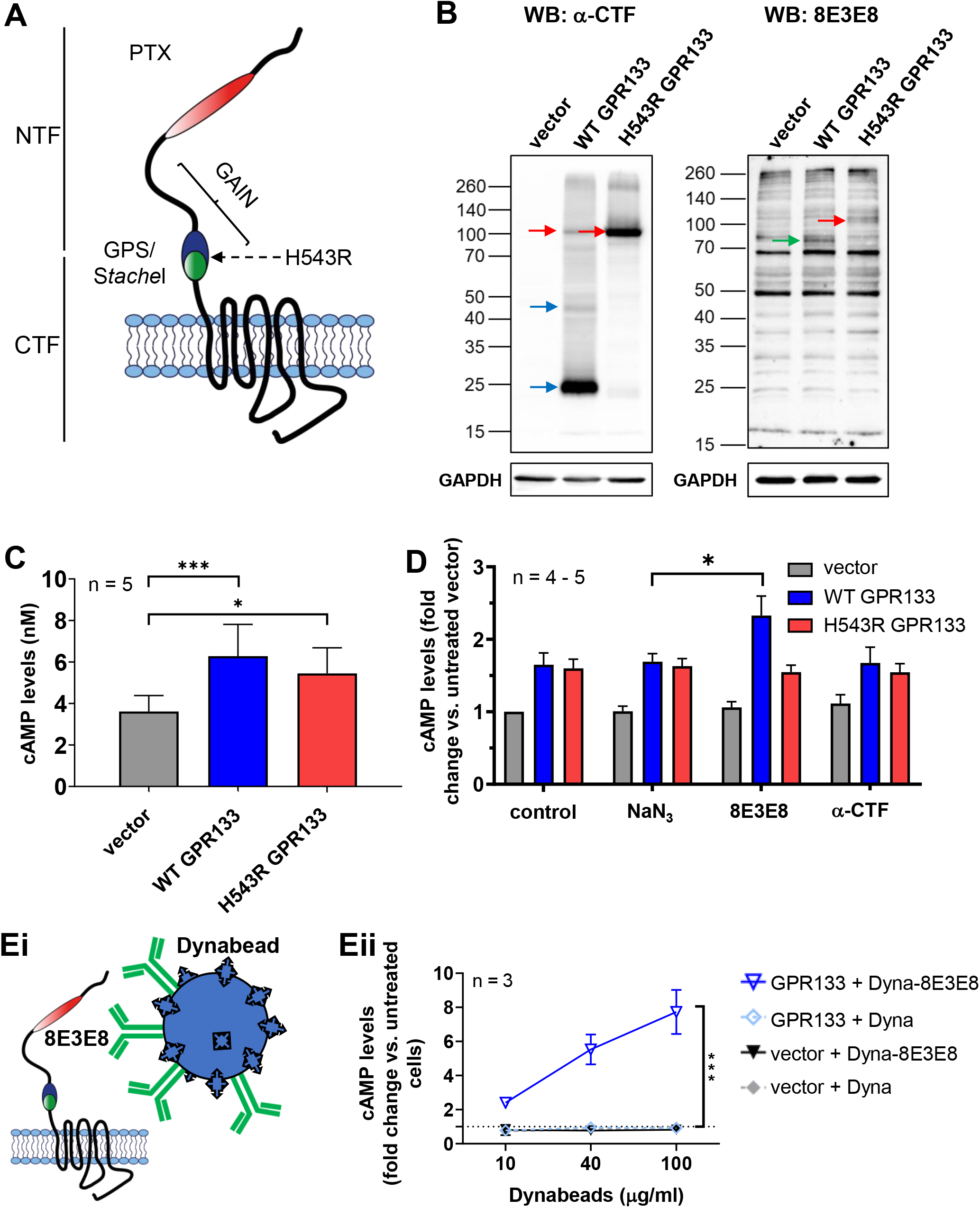
Antibody stimulation increases cAMP levels in GBM cells overexpressing GPR133. (**A**) An untagged version of GPR133 was used for these experiments and the point mutation H543R was introduced to prevent receptor cleavage. (**B**) Overexpression of GPR133 and the cleavage-deficient mutant H543R in GBML137 is shown by western blots of whole cell lysates. Upon overexpression of GPR133, the full-length receptor (red arrow), the CTF (blue arrow) and the NTF (green arrow) are detectable. When H543R is overexpressed, only a single band representing the uncleaved full-length receptor (red arrow) is detected. (**C**) cAMP levels increase significantly after overexpression of GPR133 or H543R compared to the vector control in GBML137 (F_(2,12)_=14.01, p=0.0007, one way ANOVA, Tukey’s *post hoc* test: vector vs. GPR133 *p=0.0006; vector vs. H543R *p=0.0113)). Bars represent the mean ± SEM of 5 individual experiments. (**D**) cAMP levels following treatment of GBML137 overexpressing GPR133 with different antibodies. NaN_3_ served as solvent control. Treatment with 8E3E8 leads to a significant increase of cAMP levels in cells overexpressing wild-type GPR133. The mutant H543R did not respond to the stimulus. Treating GBML137 overexpressing GPR133 or H543R with a C-terminal antibody did not change cAMP concentrations. Data points are shown as mean ± SEM of 4 – 5 individual experiments. There was a significant difference between the GPR133 + NaN_3_ and GPR133 + 8E3E8 condition (F_(2,42)_=33.66, p<0.0001, Tukey’s *post hoc* test: GPR133 + NaN_3_ and GPR133 + 8E3E8 **p=0.0091). (**Ei**) Patient-derived GBM cells (GBML137) overexpressing an un-tagged GPR133 construct were treated with Dynabeads® conjugated to 8E3E8. (**Eii**) Treatment with 8E3E8-conjugated Dynabeads® leads to a concentration-dependent increase of cAMP levels in GBML137 overexpressing GPR133, compared to the treatment with unconjugated beads. For all experiments, unconjugated Dynabeads® served as a control. Data points represent the mean ± SEM of 3 individual experiments. Among GPR133-expressing cells, there was a statistically significant difference between treatment with unconjugated Dynabeads® and Dynabeads® conjugated to 8E3E8 (F_(1,12)_=64.00, ***p<0.001, two-way ANOVA).

We also tested whether 8E3E8-coated Dynabeads® stimulate GPR133 signaling in patient-derived GBM cells (**Fig. 7Ei**), similar to their effect in HEK293T cells. Indeed, treatment of GBML137 cells overexpressing GPR133 with 8E3E8-conjugated Dynabeads® significantly increased intracellular cAMP levels, compared to cells treated with unconjugated Dynabeads® (**Fig. 7Eii**, F_(1,12)_=64.00, ***p<0.001, two way ANOVA). Overall, these findings indicate that antibody-induced GPR133 activation occurs in multiple cellular contexts, including GBM cells.

## Discussion

Our findings provide evidence that the binding of antibodies to the N-terminus of GPR133, proximal to the GAIN domain, result in receptor activation and increased canonical signaling. Following treatment of GPR133-expressing cells with these activating antibodies, we detect antibody-NTF complexes in the supernatant. Furthermore, NTF is enriched in the supernatant after treatment with the antibody in half of the trials. Finally, the effect depends on the autoproteolytic cleavage of the receptor, because an uncleavable mutant GPR133 (H543R) does not show modulation of its signaling by the same antibodies that activate the wild-type receptor. These findings suggest that the antibody-induced receptor activation is mediated by antibody-mediated dissociation of the NTF from the CTF. This mechanism is supported by our previous finding that NTF-CTF dissociation at the plasma membrane boosts receptor signaling (24), as well as the fact that GPR133 deletion mutants lacking the NTF exhibit significantly increased signaling relative to the wild-type receptor (9). Ultimately, we think that this antibody-induced NTF-CTF dissociation leads to the unveiling of the endogenous tethered agonist immediately distal to the GPS autoproteolysis site, the *Stachel* sequence, and a boost in receptor activation (**Fig. 8**).

**Figure 8:**
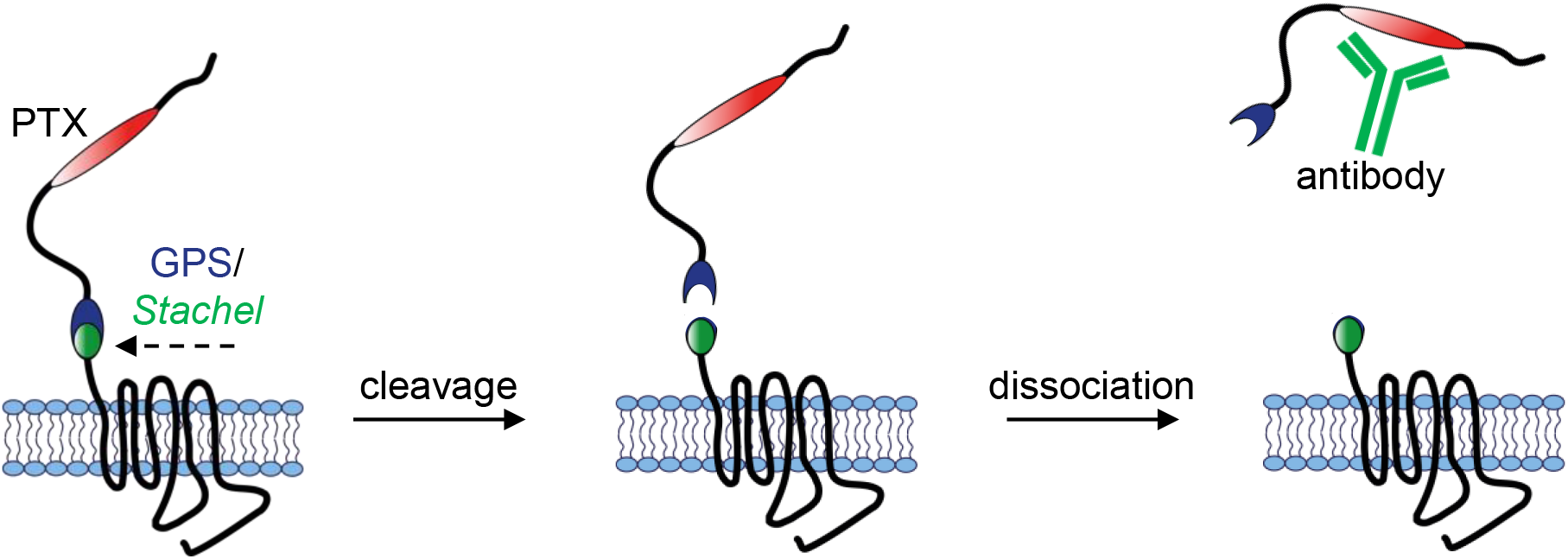
Proposed mechanism of GPR133 activation by N-terminal antibody binding. After autoproteolytic cleavage of GPR133, the NTF and CTF stay non-covalently bound to each other until antibody binding results in increased dissociation of the NTF from the CTF.

The specificity of the effect of the 8E3E8 antibody against the PTX domain of the N-terminus was demonstrated in our PTX deletion mutant GPR133. This mutant is autoproteolytically cleaved, but its basal level of signaling is reduced relative to WT GPR133. This effect may be due to conformational changes within the extracellular domain, the reduced length of the N-terminus after deleting the PTX domain, or lack of binding to PTX-specific ligands that help activate the receptor. While the PTX deletion mutant GPR133 is not modulated by 8E3E8 treatment, its signaling is still boosted by treatment with the anti-HA antibody when the receptor is HA-tagged, suggesting the underlying mechanisms behind antibody modulation are still active, even when critical domains are missing but cleavage is preserved. The fact that antibody binding to the PTX domain of the N-terminus of GPR133 increases signaling, while PTX deletion has the opposite effect, suggests a crucial function for this particular domain in receptor function, possibly through ligand binding. This hypothesis is supported by the recent identification of Plexin Domain-Containing Protein 2 (Plxdc2) as an activating ligand of GPR133, via an interaction with the PTX domain (30).

Of particular interest is the observation that coating beads with GPR133-activating antibodies produces an even more pronounced boost in receptor signaling. We theorize that this phenomenon can be explained by two different mechanisms. First, the coated beads, which in our *in vitro* culture system do not stay suspended in the culture medium but precipitate and make contact with the surface of cells, may significantly increase the effective concentration of antibody at the plasma membrane. Second, the beads, by virtue of their bulk, may represent a rigid surface for attachment of the antibodies, so that they facilitate dissociation of the antibody-bound NTF from the CTF. In essence, through their constant dynamic motions the beads may mimic the mechanical force postulated to help pull the NTF off the CTF. Third, the mechanical forces applied onto the receptor by the beads may cause conformational changes which lead to activation. Further experimentation will be required to discriminate among these mechanisms.

Previous studies showed that antibodies and other biologics directed against the extracellular domains of aGPCRs can indeed modulate receptor activation. As an example, treatment of CD97 (ADGRE5) with antibodies targeting the N-terminus outside the GAIN domain results in shedding of the receptor and formation of complexes, containing the antibody and a soluble fragment after shedding, in the supernatant of mouse splenocytes (31). Furthermore, a recent study on EMR2 (ADGRE2) showed increased canonical signaling after treatment with an antibody binding the N-terminus (32). These findings are in agreement with ours and support a shared mechanism of antibody-mediated NTF-CTF dissociation as the impetus for increased receptor activation.

Our data indicate that antibody-mediated enhancement of GPR133 signaling depends on the autoproteolytic cleavage of the receptor. This finding is consistent with our proposed model, in which the effect is mediated by NTF-CTF dissociation. However, there exist examples of other aGPCRs whose modulation by biologics does not depend on this cleavage. For example, treatment of GPR56 (ADGRG1) with monobodies targeting its N-terminus, both inside and outside the GAIN domain, modulates canonical signaling independent of cleavage (33). The discrepancy suggests activation mechanisms may vary among individual aGPCRs. Since both of our signaling-modulating antibodies target N-terminal sequences outside of the GAIN domain of GPR133, it will be interesting to test in the future whether anti-GAIN antibodies elicit effects on NTF-CTF dissociation and signaling and, if yes, the direction of modulation, activating or inhibitory.

In summary, this study provides a paradigm for the use of biologics in modulation of GPR133 signaling. Such biologics can serve as molecular tools toward better understanding receptor activation, but can also be used therapeutically in appropriate disease contexts. For example, given our discovery that GPR133 is required for GBM growth, engineering inhibitory anti-GPR133 antibodies would lay the foundation for testing effects on tumor biology (21). Alternatively, given the *de novo* expression of GPR133 in GBM relative to healthy non-neoplastic brain tissue, anti-GPR133 internalizing antibodies could be used to deliver antibody-drug conjugates to GBM cells (21,28,34).

## Experimental procedures

### Generation of GPR133 constructs

All GPR133 constructs used in this study are listed in **Table 1**. HF-GPR133 (HF = N-terminal Hemagglutinin-tag and C-terminal FLAG-tag) was available from previous studies (9,14). The H543R point mutant from HF-GPR133 was generated by site-directed mutagenesis using the Q5® Site-Directed Mutagenesis Kit (NEB, Cat# E0554S), following the manufacturer’s protocol. The PTX domain deletion construct HA-GPR133 ΔPTX was generated from HA-GPR133 using a two-fragment Gibson reaction. Twin-Strep-tagged GPR133 constructs were generated with Gibson assembly, as previously described (24). Primers for mutagenesis and Gibson cloning are listed in **Table 2**.

### Antibodies

A list of antibodies used in this study is shown in **Table 3**. For the treatment of GPR133-overexpressing cells, we used commercial anti-HA, anti-FLAG and anti-CTF antibodies or a mouse monoclonal antibody we raised against the PTX domain of GPR133 (8E3E8) (21,28). For western blot analysis we used anti-HA or 8E3E8 to detect the NTF or the full-length receptor and anti-FLAG or anti-CTF to detect the CTF or the full-length receptor. After treating cells overexpressing a Twin-Strep-tagged construct of GPR133 with 8E3E8, we used an HRP-conjugated anti-Strep antibody to detect the NTF, thus forgoing the need for a secondary antibody.

### Cell culture and transient transfection

Human embryonic kidney 293 T cells (HEK293T, Takara, Cat# 632180) were cultured in Dulbecco’s modified Eagle’s medium (DMEM, Gibco, Cat# 11965-118) supplemented with sodium pyruvate (Gibco, Cat# 11360070) and 10% fetal bovine serum (FBS; Peak Serum, Cat# PS-FB2) at 37 °C and 5% CO_2_ in humidified room air. Patient-derived GBM cells were cultured in Neurobasal medium (Gibco, Cat# 21103049) supplemented with N2 (Gibco, Cat# 17-502-049), B27 (Gibco, Cat# 12587010), non-essential amino acids (Gibco, Cat# 11140050) and GlutaMax (Gibco, Cat# 35050061) at 37°C, 5% CO_2_ and 4% O_2_ in humidified air. GBM culture medium was additionally supplemented with 20 ng/mL recombinant basic Fibroblast Growth Factor (bFGF; R&D, Cat# 233-FB-01M) and 20 ng/mL Epidermal Growth Factor (EGF; R&D, Cat# 236-EG-01M) every other day. GBM cultures were established and maintained as previously described (35). In brief, specimens were obtained from patients undergoing surgery for resection of GBM after informed consent (IRB no. 12-01130). Specimens were dissected with surgical blades and enzymatically dissociated with Accutase (Innovative Cell Technologies, Cat# AT104). Glioblastoma cells were grown as attached cultures on cell cultured dishes, pretreated with poly-L-ornithine (Sigma, Cat# P4957) and laminin (Thermo Fisher, Cat# 23017015). HEK293T cells were transfected with plasmid DNA using Lipofectamine 2000 (Invitrogen, Cat# 11668-019) and GBM cells were transfected with plasmid DNA using Lipofectamine 2000 Stem reagent (Thermo Fisher, Cat# STEM00008), following the manufacturer’s protocol.

### Homogeneous Time Resolved Fluorescence (HTRF)-based cAMP assays

Twenty-four hours after transfection, cells were seeded onto 96-well plates, pretreated with poly-L-ornithine and laminin, at a density of 75,000 cells (HEK293T) or 100,000 cells (GBM) per well. Forty-eight (HEK293T cells) to seventy-two (GBM cells) hours after transfection, we added 1 mM 3-isobutyl-1-methylxanthine (IBMX, Sigma-Aldrich, Cat# I7018-100MG) to the cells and incubated them at 37 °C for 30 – 60 min. Cells were lysed and cAMP concentrations were measured using the cAMP Gs dynamic kit (CisBio, Cat# 62AM4PEC) on the FlexStation 3 (Molecular Devices) according to the manufacturer’s protocol. For antibody stimulation experiments, antibodies (listed in **Table 3**), sodium azide (NaN_3_), which served as a solvent control because all antibodies were stored in solution containing NaN_3_, (0.015 mM, 0.06 mM, 0.15 mM) or antibody-coupled Dynabeads® (Thermo Fisher, Cat# 14311D) were added to the cells in cell culture medium one hour prior to the IBMX treatment. For antibody coupling to Dynabeads®, we followed the manufacturer’s protocol.

### Assessment of Dynabead® motion

To assess the amount of molecular motion or immobilization of Dynabeads® on monolayer cell surfaces, cells were treated with Dynabeads® as described above, followed by microscopic video capture for 200 frames over a time-course of 5 seconds. Videos were analyzed in ImageJ software by thresholding and inverting each frame to allow for specific detection of the Dynabeads® as distinct objects of correct size. Detected Dynabeads® were then tracked over 200 frames and individual bead motion tracks were plotted superimposed with the same origin to visualize the motion. Mean square displacement (MSD) was calculated as 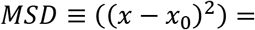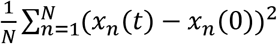.

### Enzyme-Linked Immunosorbent Assay (ELISA)

HEK293T cells, transfected with the empty vector or GPR133, were seeded onto 96-well plates as described above. Forty-eight hours after transfection (24 h after seeding), antibodies or NaN_3_ were added to the cells in cell culture medium. Cells were washed once with cold HBSS (+Ca^2+^/+Mg^2+^), fixed with 4% paraformaldehyde (PFA, Sigma-Aldrich, Cat# P6148) for 20 min at room temperature (RT) and blocked with cell culture medium containing 10% FBS for 1 h at RT. Cells were then incubated with horseradish peroxidase (HRP)-conjugated secondary antibodies (1:1000, chicken-anti mouse IgG Invitrogen Cat# A15975 or chicken-anti rabbit IgG Invitrogen Cat# A15987, Table 4) in cell culture medium containing 10% FBS for 1 h at RT. After 3 washes with phosphate-buffered saline (PBS), cells were incubated with TMB (3,3’, 5,5’-tetramethylbenzidine)-stabilized chromogen (Thermo Fisher, Cat# SB02) for 15 min. The reaction was stopped by adding an equal volume of acidic stop solution (Thermo Fisher, Cat# SS04) and optical density was measured at 450 nm.

### Western blot analysis of whole cell lysates

Cells were washed with PBS and lysed in RIPA buffer (Thermo, Cat#89900) supplemented with Halt protease inhibitor cocktail (Thermo, Cat# 78429) and 1% n-dodecyl β-D-maltoside (DDM) (Thermo, Cat# BN2005). Lysates were incubated for 15 min on ice, sonicated in a water-bath Bioruptor (Diagenode, Cat# UCD-300) and precleared by centrifugation at 15,000 x g for 10 min at 4 °C. Protein concentrations were measured using the DC protein assay kit II (BioRad, Cat# 5000112). Laemmli buffer (BioRad, Cat# 1610747), supplemented with β-mercaptoethanol, was then added and lysates were incubated at 37 °C for 30 min. Twenty μg of protein were separated by SDS-PAGE and transferred to 0.2 μm nitrocellulose membranes (BioRad, Cat# 1620112). Membranes were blocked in 2% bovine serum albumin (BSA) in TBS-Tween for 1 h at RT and incubated with primary antibodies (listed in **Table 3**) at 4 °C overnight. Following incubation with Alexa Fluor or HRP-conjugated secondary antibodies (listed in **Table 4**) for 1 h at RT, images were acquired using the iBrightFL1000 system (Invitrogen). Signals were detected by fluorescence or chemiluminescence (Thermo Scientific, Cat# 34577), depending on the secondary antibody. Densitometric quantification of band intensities was carried out using ImageJ.

### Affinity purification of Strep-tagged GPR133

Twin-Strep-tagged GPR133 (**Table 1**) was affinity-purified from culture supernatants using Strep-Tactin® XT coated magnetic beads (MagStrep “type3” XT Beads, IBA, Cat# 2-4090-002), according to the manufacturer’s protocol. In brief, HEK293T cells were transfected with N-terminally TwinStrep-tagged constructs of wild-type (WT) GPR133, as described above. Twenty-four hours after transfection, cell culture medium was replaced by DMEM supplemented with 2% FBS. Seventy-two hours after transfection, cells were treated with 1.8 μg/ml 8E3E8 antibody for 1 – 4 h at 37 °C. Whole cell lysates were prepared as described above. Culture supernatant was treated with 10X Buffer W (IBA, Cat# 2-1003-100) and BioLock (IBA, Cat# 2-0205-250) for 15 minutes on ice and precleared by centrifugation at 15,000 x g for 15 min at 4 °C. The supernatant was incubated with MagStrep “type3” XT Beads overnight at 4°C. The next day, beads were collected with a magnetic separator and washed 3 times with 1x Buffer W. Proteins were eluted in two consecutive elutions with 1X biotin elution buffer BXT (IBA, Cat# 2-1042-025). After pooling the elutions, Laemmli buffer with β-mercaptoethanol was added and proteins were analyzed by western blot as described above.

### Deglycosylation

Whole cell lysates or eluted proteins after affinity purification were treated with Protein Deglycosylation Mix II (NEB, Cat# P6044), according to the manufacturer’s protocol. Deglycosylation was performed under denaturing conditions for 16 hours at 37 °C. Samples were stored in Laemmli buffer, supplemented with β-mercaptoethanol, and analyzed by western blot.

### Immunofluorescent staining

HEK293T cells were transfected and cultured on dishes coated with poly-L-ornithine and laminin, as described above. Cells were fixed with 4% PFA for 30 min at RT, followed by block and permeabilization with 10% BSA in PBS supplemented with 0.1% Triton X-100 for 1 h at RT. Cells were then incubated with a primary anti-HA antibody (Sigma, Cat# H3663) in 1% BSA in PBS + 0.1% Triton X-100 at 4 °C overnight. The next day, cells were washed with PBS + 0.1% Triton X-100 and stained with donkey anti-mouse AlexaPlus 488 IgG (H+L) secondary antibody for 1 h at RT. Nuclei were counterstained with 500 ng/mL 4′,6-diamidino-2-phenylindole (DAPI) for 10 min at RT. Microscopy was conducted on a Zeiss Axiovert epifluorescent microscope.

### Statistical analysis

Statistical analysis was performed using GraphPad Prism (version 8.4.3). Population statistics are represented as mean ± standard error of the mean (SEM) throughout. Statistical significance was calculated using either Students t-test, one-way analysis of variance (ANOVA), or two-way ANOVA with Tukey’s *post hoc* test for multiple comparisons. P values <0.05 were considered statistically significant (*p<0.05; **p<0.01; ***p<0.001).

## Data availability

All data are contained within the manuscript.

## Supporting information

The article contains supporting information.

## Author contributions

Conceived and designed the experiments: GS, IL, DGP. Performed the experiments: GS, JDF. Analyzed the data: GS, JDF. Wrote and edited the manuscript: GS, JDF, IL, DGP.

## References

1. Krishnan, A., Nijmeijer, S., de Graaf, C., and Schiöth, H. B. (2016) Classification, Nomenclature, and Structural Aspects of Adhesion GPCRs. Handbook of experimental pharmacology 234, 15–41

2. Hamann, J., Aust, G., Araç, D., Engel, F. B., Formstone, C., Fredriksson, R., Hall, R. A., Harty, B. L., Kirchhoff, C., Knapp, B., Krishnan, A., Liebscher, I., Lin, H. H., Martinelli, D. C., Monk, K. R., Peeters, M. C., Piao, X., Prömel, S., Schöneberg, T., Schwartz, T. W., Singer, K., Stacey, M., Ushkaryov, Y. A., Vallon, M., Wolfrum, U., Wright, M. W., Xu, L., Langenhan, T., and Schiöth, H. B. (2015) International Union of Basic and Clinical Pharmacology. XCIV. Adhesion G protein-coupled receptors. Pharmacological reviews 67, 338–367

3. Langenhan, T. (2020) Adhesion G protein-coupled receptors-Candidate metabotropic mechanosensors and novel drug targets. Basic & clinical pharmacology & toxicology 126 Suppl 6, 5–16

4. Kaczmarek, I., Suchý, T., Prömel, S., Schöneberg, T., Liebscher, I., and Thor, D. (2021) The relevance of adhesion G protein-coupled receptors in metabolic functions. Biological chemistry

5. Gad, A. A., and Balenga, N. (2020) The Emerging Role of Adhesion GPCRs in Cancer. ACS pharmacology & translational science 3, 29–42

6. Scholz, N., Langenhan, T., and Schöneberg, T. (2019) Revisiting the classification of adhesion GPCRs. Annals of the New York Academy of Sciences 1456, 80–95

7. Hayflick, J. S. (2000) A family of heptahelical receptors with adhesion-like domains: a marriage between two super families. Journal of receptor and signal transduction research 20, 119–131

8. Araç, D., Boucard, A. A., Bolliger, M. F., Nguyen, J., Soltis, S. M., Südhof, T. C., and Brunger, A. T. (2012) A novel evolutionarily conserved domain of cell-adhesion GPCRs mediates autoproteolysis. The EMBO journal 31, 1364–1378

9. Liebscher, I., Schön, J., Petersen, S. C., Fischer, L., Auerbach, N., Demberg, L. M., Mogha, A., Cöster, M., Simon, K. U., Rothemund, S., Monk, K. R., and Schöneberg, T. (2015) A Tethered Agonist within the Ectodomain Activates the Adhesion G Protein-Coupled Receptors GPR126 and GPR133. Cell reports 10, 1021

10. Stoveken, H. M., Hajduczok, A. G., Xu, L., and Tall, G. G. (2015) Adhesion G protein-coupled receptors are activated by exposure of a cryptic tethered agonist. Proceedings of the National Academy of Sciences of the United States of America 112, 6194–6199

11. Petersen, S. C., Luo, R., Liebscher, I., Giera, S., Jeong, S. J., Mogha, A., Ghidinelli, M., Feltri, M. L., Schöneberg, T., Piao, X., and Monk, K. R. (2015) The adhesion GPCR GPR126 has distinct, domain-dependent functions in Schwann cell development mediated by interaction with laminin-211. Neuron 85, 755–769

12. Monk, K. R., Hamann, J., Langenhan, T., Nijmeijer, S., Schöneberg, T., and Liebscher, I. (2015) Adhesion G Protein-Coupled Receptors: From In Vitro Pharmacology to In Vivo Mechanisms. Molecular pharmacology 88, 617–623

13. Demberg, L. M., Rothemund, S., Schöneberg, T., and Liebscher, I. (2015) Identification of the tethered peptide agonist of the adhesion G protein-coupled receptor GPR64/ADGRG2. Biochemical and biophysical research communications 464, 743–747

14. Demberg, L. M., Winkler, J., Wilde, C., Simon, K. U., Schön, J., Rothemund, S., Schöneberg, T., Prömel, S., and Liebscher, I. (2017) Activation of Adhesion G Protein-coupled Receptors: AGONIST SPECIFICITY OF STACHEL SEQUENCE-DERIVED PEPTIDES. The Journal of biological chemistry 292, 4383–4394

15. Müller, A., Winkler, J., Fiedler, F., Sastradihardja, T., Binder, C., Schnabel, R., Kungel, J., Rothemund, S., Hennig, C., Schöneberg, T., and Prömel, S. (2015) Oriented Cell Division in the C. elegans Embryo Is Coordinated by G-Protein Signaling Dependent on the Adhesion GPCR LAT-1. PLoS genetics 11, e1005624

16. Kishore, A., Purcell, R. H., Nassiri-Toosi, Z., and Hall, R. A. (2016) Stalk-dependent and Stalk-independent Signaling by the Adhesion G Protein-coupled Receptors GPR56 (ADGRG1) and BAI1 (ADGRB1). The Journal of biological chemistry 291, 3385–3394

17. Wilde, C., Fischer, L., Lede, V., Kirchberger, J., Rothemund, S., Schöneberg, T., and Liebscher, I. (2016) The constitutive activity of the adhesion GPCR GPR114/ADGRG5 is mediated by its tethered agonist. FASEB journal : official publication of the Federation of American Societies for Experimental Biology 30, 666–673

18. Scholz, N., Guan, C., Nieberler, M., Grotemeyer, A., Maiellaro, I., Gao, S., Beck, S., Pawlak, M., Sauer, M., Asan, E., Rothemund, S., Winkler, J., Prömel, S., Nagel, G., Langenhan, T., and Kittel, R. J. (2017) Mechano-dependent signaling by Latrophilin/CIRL quenches cAMP in proprioceptive neurons. eLife 6

19. Favara, D. M., Liebscher, I., Jazayeri, A., Nambiar, M., Sheldon, H., Banham, A. H., and Harris, A. L. (2021) Elevated expression of the adhesion GPCR ADGRL4/ELTD1 promotes endothelial sprouting angiogenesis without activating canonical GPCR signalling. Scientific reports 11, 8870

20. Vizurraga, A., Adhikari, R., Yeung, J., Yu, M., and Tall, G. G. (2020) Mechanisms of adhesion G protein-coupled receptor activation. The Journal of biological chemistry 295, 14065–14083

21. Bayin, N. S., Frenster, J. D., Kane, J. R., Rubenstein, J., Modrek, A. S., Baitalmal, R., Dolgalev, I., Rudzenski, K., Scarabottolo, L., Crespi, D., Redaelli, L., Snuderl, M., Golfinos, J. G., Doyle, W., Pacione, D., Parker, E. C., Chi, A. S., Heguy, A., MacNeil, D. J., Shohdy, N., Zagzag, D., and Placantonakis, D. G. (2016) GPR133 (ADGRD1), an adhesion G-protein-coupled receptor, is necessary for glioblastoma growth. Oncogenesis 5, e263

22. Frenster, J. D., Inocencio, J. F., Xu, Z., Dhaliwal, J., Alghamdi, A., Zagzag, D., Bayin, N. S., and Placantonakis, D. G. (2017) GPR133 Promotes Glioblastoma Growth in Hypoxia. Neurosurgery 64, 177–181

23. Dolecek, T. A., Propp, J. M., Stroup, N. E., and Kruchko, C. (2012) CBTRUS statistical report: primary brain and central nervous system tumors diagnosed in the United States in 2005-2009. Neuro-oncology 14 Suppl 5, v1–49

24. Frenster, J. D., Stephan, G., Ravn-Boess, N., Bready, D., Wilcox, J., Kieslich, B., Wilde, C., Sträter, N., Wiggin, G. R., Liebscher, I., Schöneberg, T., and Placantonakis, D. G. (2021) Functional impact of intramolecular cleavage and dissociation of adhesion G protein-coupled receptor GPR133 (ADGRD1) on canonical signaling. The Journal of biological chemistry, 100798

25. Bohnekamp, J., and Schöneberg, T. (2011) Cell adhesion receptor GPR133 couples to Gs protein. The Journal of biological chemistry 286, 41912–41916

26. Gupte, J., Swaminath, G., Danao, J., Tian, H., Li, Y., and Wu, X. (2012) Signaling property study of adhesion G-protein-coupled receptors. FEBS letters 586, 1214–1219

27. Fischer, L., Wilde, C., Schöneberg, T., and Liebscher, I. (2016) Functional relevance of naturally occurring mutations in adhesion G protein-coupled receptor ADGRD1 (GPR133). BMC genomics 17, 609

28. Frenster, J. D., Kader, M., Kamen, S., Sun, J., Chiriboga, L., Serrano, J., Bready, D., Golub, D., Ravn-Boess, N., Stephan, G., Chi, A. S., Kurz, S. C., Jain, R., Park, C. Y., Fenyo, D., Liebscher, I., Schöneberg, T., Wiggin, G., Newman, R., Barnes, M., Dickson, J. K., MacNeil, D. J., Huang, X., Shohdy, N., Snuderl, M., Zagzag, D., and Placantonakis, D. G. (2020) Expression profiling of the adhesion G protein-coupled receptor GPR133 (ADGRD1) in glioma subtypes. Neuro-oncology advances 2, vdaa053

29. Beliu, G., Altrichter, S., Guixà-González, R., Hemberger, M., Brauer, I., Dahse, A. K., Scholz, N., Wieduwild, R., Kuhlemann, A., Batebi, H., Seufert, F., Pérez-Hernández, G., Hildebrand, P. W., Sauer, M., and Langenhan, T. (2021) Tethered agonist exposure in intact adhesion/class B2 GPCRs through intrinsic structural flexibility of the GAIN domain. Molecular cell 81, 905–921.e905

30. Bianchi, E., Sun, Y., Almansa-Ordonez, A., Woods, M., Goulding, D., Martinez-Martin, N., and Wright, G. J. (2021) Control of oviductal fluid flow by the G-protein coupled receptor Adgrd1 is essential for murine embryo transit. Nature communications 12, 1251

31. de Groot, D. M., Vogel, G., Dulos, J., Teeuwen, L., Stebbins, K., Hamann, J., Owens, B. M., van Eenennaam, H., Bos, E., and Boots, A. M. (2009) Therapeutic antibody targeting of CD97 in experimental arthritis: the role of antigen expression, shedding, and internalization on the pharmacokinetics of anti-CD97 monoclonal antibody 1B2. Journal of immunology (Baltimore, Md. : 1950) 183, 4127–4134

32. Bhudia, N., Desai, S., King, N., Ancellin, N., Grillot, D., Barnes, A. A., and Dowell, S. J. (2020) G Protein-Coupling of Adhesion GPCRs ADGRE2/EMR2 and ADGRE5/CD97, and Activation of G Protein Signalling by an Anti-EMR2 Antibody. Scientific reports 10, 1004

33. Salzman, G. S., Zhang, S., Gupta, A., Koide, A., Koide, S., and Araç, D. (2017) Stachel-independent modulation of GPR56/ADGRG1 signaling by synthetic ligands directed to its extracellular region. Proceedings of the National Academy of Sciences of the United States of America 114, 10095–10100

34. Stephan, G., Ravn-Boess, N., and Placantonakis, D. G. (2021) Adhesion G protein-coupled receptors in glioblastoma. Neuro-oncology advances 3, vdab046

35. Frenster, J. D., and Placantonakis, D. G. (2018) Establishing Primary Human Glioblastoma Tumorsphere Cultures from Operative Specimens. Methods in molecular biology (Clifton, N.J.) 1741, 63–69

