## Supporting Information for "Activation of the adhesion GPCR GPR133 (ADGRD1) by antibodies targeting the N-terminus"

### Tables

Table 1: GPR133 constructs

| name | backbone | tags | cleavage |
|---|---|---|---|
| GPR133 (HF-GPR133) | pcDps | HA (N-terminal)<br>FLAG (C-terminal) | yes |
| H543R (HF-GPR133 (H543R)) | pcDps | HA (N-terminal)<br>FLAG (C-terminal) | no |
| GPR133 (codon-optimized) | pLVX | - | yes |
| H543R (codon-optimized) | pLVX | - | no |
| HA-GPR133 | pLVX | HA (N-terminal) | yes |
| HA-GPR133 ΔPTX | pLVX | HA (N-terminal) | yes |
| Strep-GPR133 | pLVX | Twin-Strep (N-terminal) | yes |
| Strep-GPR133 (H543R) | pLVX | Twin-Strep (N-terminal) | no |

Table 2: Primer sequences

| primer | sequence |
|---|---|
| HF-GPR133 (H543R) for | CGCTGCACTCGACTCACCAACTTTG |
| HF-GPR133 (H543R) rev | GCAGACGGAGTAGGTGAG |
| ΔPTX Gibson Fragment 1 for | GCATTTACCTGAAAGAGGAACATCCCATCATAACCAACCTGACAG |
| ΔPTX Gibson Fragment 1 rev | GGTTAGCTCCTTCGGTCCTCC |
| ΔPTX Gibson Fragment 2 for | GATAACACTGCGGCCAACTTACTTC |
| ΔPTX Gibson Fragment 2 rev | AGGTTGGTTATGATGGGATGTTCCTCTTTCAGGTAAATGCCTTTG |

Table 3: Primary antibodies

| antibody | target | species | Company | Cat# |
|---|---|---|---|---|
| anti-HA | HA-tag | mouse | Sigma | H3663-100UL |
| anti-FLAG | FLAG-tag | rabbit | Sigma | F3165-.2MG |
| 8E3E8 | GPR133 NTF | mouse | Genscript | - |
| anti-CTF | GPR133 CTF | rabbit | Sigma | HPA042395 |
| anti-Strep-HRP | Strep-tag | mouse | IBA | 2-1509-001 |
| anti-GAPDH | GAPDH | goat | Invitrogen | PA1-9046 |

Table 4: Secondary antibodies

| antibody | detection | species | Company | Cat# |
|---|---|---|---|---|
| anti-mouse IgG | Alexa Fluor plus 488 | donkey | Invitrogen | A32766 |
| anti-rabbit IgG | Alexa Fluor plus 555 | donkey | Invitrogen | A32794 |
| anti-goat IgG | Alexa Fluor plus 647 | donkey | Invitrogen | A32849 |
| anti-mouse IgG | HRP / chemiluminescence | chicken | Invitrogen | A15975 |
| anti-rabbit IgG | HRP / chemiluminescence | chicken | Invitrogen | A15987 |

### Supplementary Figures

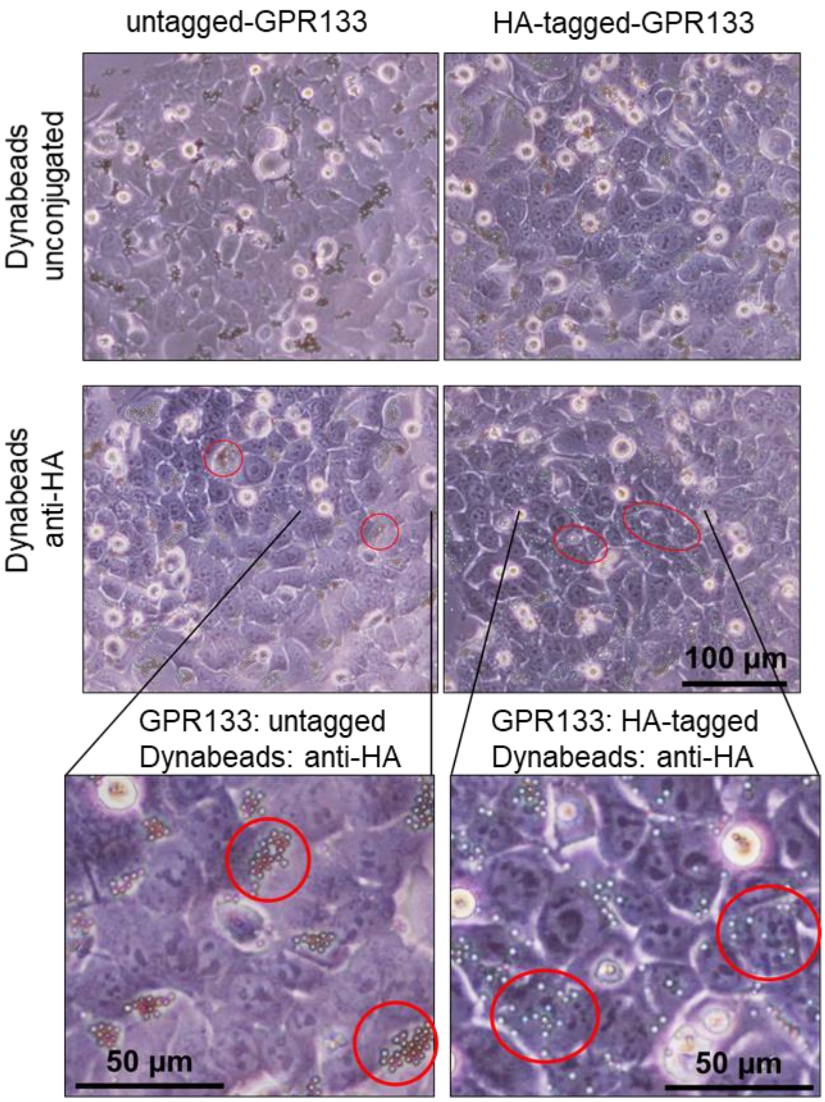


**Supplementary Figure 1: Anti-HA-conjugated Dynabeads® adhere to the plasma membrane of HEK293T cells overexpressing HA-tagged GPR133.** Representative micrographs of HEK293T cells treated with Dynabeads®. Unconjugated Dynabeads® (top panels) or anti-HA-conjugated Dynabeads® (bottom panels) were incubated with cells overexpressing untagged GPR133 (left panel) or HA-tagged GPR133 (right panel). When anti-HA-conjugated Dynabeads® and HA-tagged GPR133 are present, beads spread out across cell surfaces. In all other conditions, beads cluster between cells. Red circles highlight examples of unbound clustered Dynabeads® (left), as well as cell surface-bound anti-HA-conjugated Dynabeads® (right).

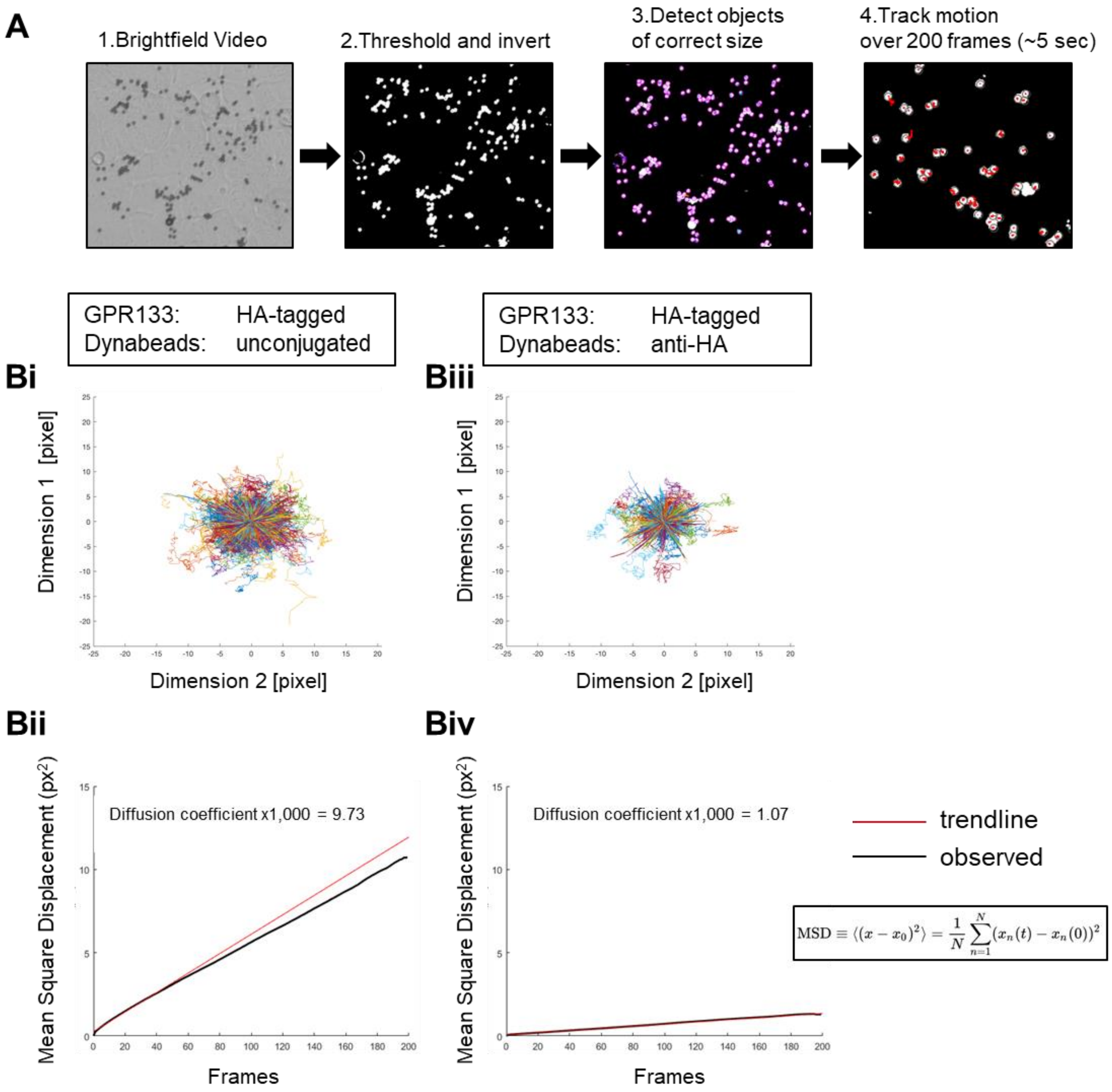


**Supplementary Figure 2: Anti-HA-conjugated Dynabeads® are immobilized upon binding to HEK293T cells overexpressing HA-tagged GPR133.** Diffusivity of Dynabeads® on cell surfaces was assessed by microscopic video capture and motility tracing in ImageJ. HEK293T cells overexpressing HA-tagged GPR133 were treated with either unconjugated Dynabeads® or anti-HA-conjugated Dynabeads®. (**A**) Five second-long brightfield videos consisting of 200 frames were captured under the microscope and analyzed in ImageJ. Each frame was thresholded and inverted, and beads were automatically identified as round objects of the correct size. The motion was tracked through all frames as the displacement of the center of each detected object, creating a motion path for each Dynabead®. (**Bi, Biii**) The motion paths of all Dynabeads® within each condition were superimposed with the same origin and (**Bii, Biv**) the mean square displacement of all beads within each experimental condition was plotted as a function of time (frames). Anti-HA-conjugated Dynabeads® were relatively immobile on HA-GPR133-expressing HEK293T cells when compared to unconjugated Dynabeads® (Diffusion coefficient x1,000 of 1.07 and 9.73 respectively).

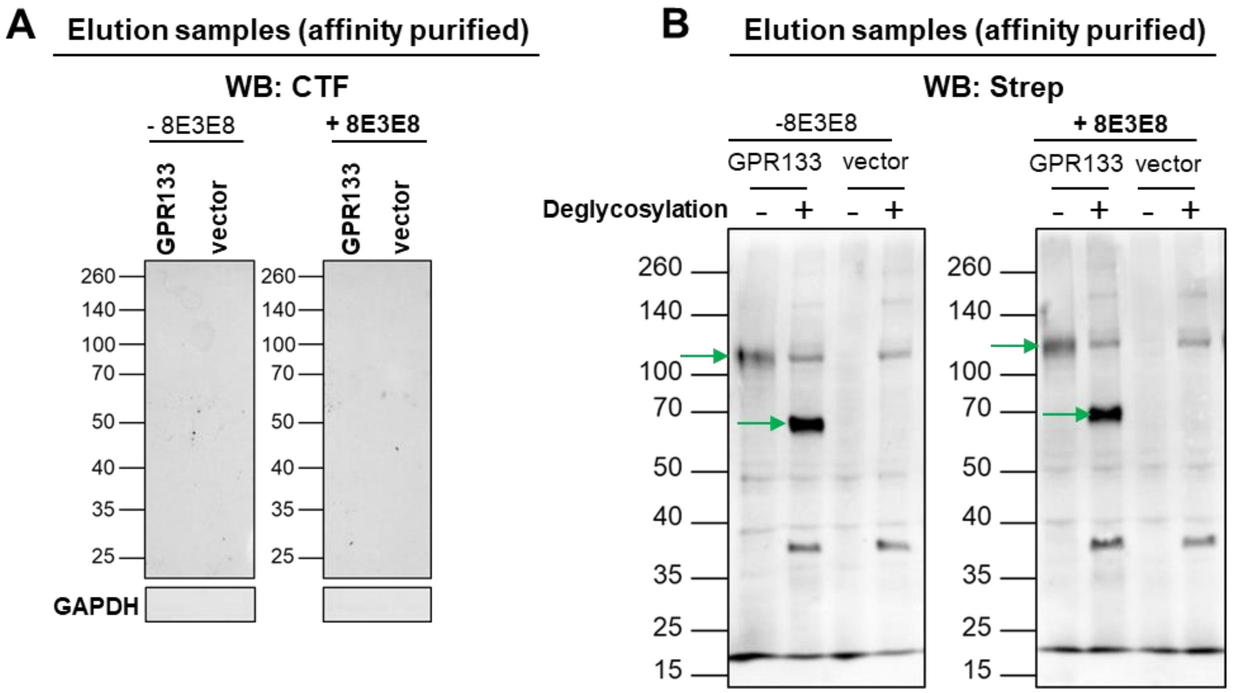


**Supplementary Figure 3: Quality controls in eluates.** HEK293T cells overexpressing GPR133 were treated with the 8E3E8 antibody, followed by Streptactin® purification of the supernatant, and analysis of eluates by Western blot. (**A**) Probing with anti-CTF antibody shows no signal. (**B**) Deglycosylation of elution samples following treatment with 8E3E8. The blot was stained with the anti-Strep antibody. Green arrows point to the NTF.
